# Proteomic phenotyping of stimulated Müller cells uncovers profound pro-inflammatory signaling and antigen-presenting capacity

**DOI:** 10.1101/2021.08.29.458112

**Authors:** Adrian Schmalen, Lea Lorenz, Antje Grosche, Diana Pauly, Cornelia A. Deeg, Stefanie M. Hauck

## Abstract

Müller cells are the main macroglial cells of the retina exerting a wealth of functions to maintain retinal homoeostasis. Upon pathological changes in the retina, they become gliotic with both protective and detrimental consequences. Accumulating data also provide evidence for a pivotal role of Müller cells in the pathogenesis of diabetic retinopathy (DR). While microglial cells, the resident immune cells of the retina are considered as main players in inflammatory processes associated with DR, the implication of activated Müller cells in chronic retinal inflammation remains to be elucidated. In order to assess the signaling capacity of Müller cells and their role in retinal inflammation, we performed in-depth proteomic analysis of Müller cell proteomes and secretomes after stimulation with INFγ, TNFα, IL-4, IL-6, IL-10, VEGF, TGFβ1, TGFβ2 and TGFβ3. We used both, primary porcine Müller cells and the human Müller cell line MIO-M1 for our hypothesis generating approach. Our results point towards an intense signaling capacity of Müller cells, which reacted in a highly discriminating manner upon treatment with different cytokines. Stimulation of Müller cells resulted in a primarily pro-inflammatory phenotype with secretion of cytokines and components of the complement system. Furthermore, we observed evidence for mitochondrial dysfunction, implying oxidative stress after treatment with the various cytokines. Finally, both MIO-M1 cells and primary porcine Müller cells showed several characteristics of atypical antigen-presenting cells, as they are capable of inducing MHC class I and MHC class II with co-stimulatory molecules. In line with this, they express proteins associated with formation and maturation of phagosomes. Thus, our findings underline the importance of Müller cell signaling in the inflamed retina, indicating an active role in chronic retinal inflammation underlying the pathogenesis of diabetic retinopathy.

## 1 Introduction

Neurodegenerative diseases of the retina are characterized by progressive photoreceptor damage eventually resulting in vision loss (1). Accumulating evidence over the last years led to the recognition of chronic inflammation as an important part of the pathogenesis underlying this heterogeneous group of retinal diseases (2–4). While the loss of photoreceptors in inherited retinal diseases such as retinitis pigmentosa originates in various genetic mutations (5), the driving force that leads to retinal degeneration and blindness in diabetic retinopathy (DR) is a disturbed metabolism with hyperglycemia and dyslipidemia, resulting in microvascular damage (6). DR is among the most frequent causes of blindness worldwide with a rising prevalence (7). Chronic hyperglycemia in diabetes patients induces the activation of leukocytes in the periphery (8) as well as micro- and macroglial cells in the retina (9–11). This results in the release of pro-inflammatory cytokines and eventually leads to photoreceptor cell death (12, 13). Microglial cells, the resident immune cells of the retina, are acknowledged as the main drivers of retinal immune responses (14). However, growing evidence suggests that the interaction of micro- and macroglial cells essentially shapes retinal inflammation and photoreceptor degeneration (15, 16).

Retinal Müller glial cells constitute the primary macroglial cells of the retina (17). They span the entire width of the retina and are in close contact with the vitreous, the retinal blood-vessels and with all retinal neurons (18). While they maintain retinal homeostasis during steady-state conditions, activation of Müller cells under pathological conditions results in gliosis, a cellular attempt to restore insulted tissue, with both protective and detrimental effects (19). It is known that Müller cells are an important source of neurotrophic factors but also of pro-inflammatory and angiogenetic cytokines (20–23). They play an important role in DR pathogenesis (24, 25) and it was also suggested that they are involved in retinal immune responses (26–30). However, the impact of their protective or detrimental effects on retinal inflammation in DR and other neurodegenerative retinal diseases remains elusive so far. Thus, we performed an in-depth-analysis of the Müller cell proteome and secretome after stimulation with various pro- and anti-inflammatory cytokines as well as growth factors. For our analysis, we used cells of the human Müller glia cell-line MIO-M1 as well as primary porcine retinal Müller Glia (pRMG), as porcine eyes resemble human eyes regarding their anatomy (31). Furthermore, the pig is established as a useful model for research in diabetic retinopathy (32, 33). Comparative quantitative proteomic analysis allows to elucidate key proteins or pathways involved in disease pathogenesis. In addition, this approach enables the discovery of new biomarkers and has therefore proven as a valuable tool for deciphering pathophysiological key mechanisms (34, 35). Our hypothesis-generating study yielded comprehensive data on the capacity of Müller cells to react in a very differentiated manner to varying stimulants. In addition to the secretion of pro-inflammatory cytokines, we observed expression of MHC class I and II molecules and proteins that are associated with the processing of antigens. We therefore propose that Müller cells are critical modulators of the retinal immune response and might exert an antigen-presenting function. Thus, attention should be paid to their implication in chronic inflammation underlying degenerative retinal diseases.

## 2 Materials and methods

### 2.1 Cell preparation and culture

Cells were maintained in Dulbecco’s modified eagle medium (DMEM) containing 10% (*v/v*) inactivated fetal calf serum (FCS), 100 U/mL penicillin and 100 μg/mL streptomycin unless otherwise stated. Cell culture media and reagents were purchased from Gibco (Life Technologies GmbH, Darmstadt, Germany).

Ten porcine eyes from healthy pigs were kindly provided from a local abattoir. The use of porcine material from the abattoir was approved for purposes of scientific research by the appropriate board of the veterinary inspection office, Munich, Germany (registration number DE 09 162 0008-21). No experimental animals were involved in this study. Within 2h after enucleation, eyes were processed under a laminar flow hood under sterile conditions as previously described (30, 36). In short, periocular tissue was removed and the eyeballs were rinsed in 80% ethanol followed by washing with cold PBS. Afterwards, eyeballs were stored in DMEM until further processing. The eyeballs were opened circumferentially parallel to the limbus corneae, and anterior parts of the eyes were discarded. The retina was detached from the posterior eyeballs and transferred into a petri dish containing DMEM. After removal of vitreous and pigment epithelium residues, major blood-vessels were excised and the remaining retinal tissue was cut into very small fragments using micro-scissors. Resulting fragments were washed in Ringer’s solution followed by enzymatic digestion at 37 °C with papain previously activated by incubation with 1.1 μM EDTA, 0.067 μM mercaptoethanol and 5.5 μM cysteine-HCl. Enzymatic digestion was stopped after 12 min by adding serum-containing DMEM, followed by addition of Desoxyribonuclease I (Sigma-Aldrich Chemie GmbH, Taufkirchen, Germany) and trituration. After sedimentation of the cells, the supernatant was carefully removed using Pasteur pipettes. The remaining pellets were resuspended in DMEM, pooled and seeded into 6-well plates (Sarstedt, Nümbrecht, Germany). The following day, thorough panning of the plates and removal of the supernatant were performed in order to eliminate non-attached cells, yielding pure Müller cell cultures as previously described (37, 38). Cells were cultured at 37 °C and 5% CO_2_ with regular exchange of medium and repeated microscopic control of cell density and purity according to previous reports (37, 39).

The human Müller cell line Moorfields/Institute of Ophthalmology-Müller 1 (MIO-M1; RRID:CVCL_0433) was a kind gift of G. A. Limb (39). They were tested negative for mycoplasma contamination. Two days before treatment, 1×10^5^ MIO-M1 cells per well were seeded in 6-well plates and incubated at 37 °C and 5% CO_2_ until further processing.

### 2.2 Cell stimulation

The human cytokines Interleukin 10 (IL-10), Transforming Growth Factor beta-1 (TGFβ1), Transforming Growth Factor beta-2 (TGFβ2), Transforming Growth Factor beta-3 (TGFβ3), and Tumor Necrosis Factor Alpha (TNFα) were purchased from Sigma-Aldrich, Interleukin 4 (IL-4) and Interferon Gamma (IFNy) from R&D Systems/Bio-Techne (R&D Systems/Bio-Techne, Wiesbaden-Nordenstadt, Germany), and Interleukin 6 (IL-6) and Vascular Endothelial Growth Factor165 (VEGF) from PeproTech (PeproTech, Winterhude, Germany). Porcine TGFβ3 was purchased from Biozol (Biozol, Eching, Germany), whereas porcine TGFβ1, TGFβ2, IL-4, IL-6, IL-10, IFNy and TNFα were from R&D System/Bio-Techne. Since there was no porcine VEGF available, the above mentioned human VEGF was also used for stimulation of pRMG.

To diminish the influence of cytokines present in FCS, both confluent pRMG and MIO-M1 cells were rinsed two times with prewarmed serum-free medium, followed by starvation for 1 h at 37 °C and 5% CO_2_ with serum-deprived medium. Afterwards, cells were treated over night with IFNy, IL-4, IL-6, IL-10, TGFβ1, TGFβ2, TGFβ3, TNFα or VEGF_165_, respectively, in a randomized plate design at a concentration of 5 ng/mL in medium without FCS. Untreated cells cultured in serum-free medium served as a control.

### 2.3 Sample collection and proteolysis

Supernatants were collected 24 h after treatment, passed through medium equilibrated 0.2 μm Millex-GP filter units (Merck Chemicals GmbH, Darmstadt, Germany), and transferred into 2 mL Lo-Bind tubes (Eppendorf AG, Hamburg, Germany). Afterwards, cells were washed once with DPBS. 200 μL RIPA buffer containing Roche cOmplete Mini Protease Inhibitor Cocktail (Merck Chemicals GmbH) was applied directly into each well and cells were detached with a cell scraper. Lysates were transferred into freshly prepared 1.5 mL Lo-Bind tubes (Eppendorf AG). Protein concentration of the lysates was determined by Pierce BCA assay (Thermo Fisher Scientific). Ten μg protein per lysate or 400 μL supernatant per sample were digested with Lys-C and trypsin using a modified FASP procedure as described elsewhere (40, 41).

### 2.4 LC-MS/MS and quantitative analysis

LC-MSMS analysis was performed on a QExactive HF mass spectrometer (Thermo Fisher Scientific) online coupled to a UItimate 3000 RSLC nano-HPLC (Dionex, Sunnyvale, USA). Samples were automatically injected and loaded onto a C18 trap column for 5 min. Afterwards, samples were eluted and separated on a C18 analytical column (Acquity UPLC M-Class HSS T3 Column, 1.8 μm, 75 μm × 250 mm; Waters, Milford, USA). Samples were separated by a 95 min non-linear acetonitrile gradient at a flow rate of 250 nl/min. Resolution of the MS spectra was recorded at 60,000 with an AGC target of 3×10^6^ and a maximum injection time of 50 ms from 300 to 1,500 m/z. The 10 most abundant peptide ions were selected from the MS scan and fragmented via HCD. Thereby, the normalized collision energy was 27 with an isolation window of 1.6 m/z, and a dynamic exclusion of 30 s. MS/MS spectra were recorded at a resolution of 15,000 with an AGC target of 10^5^ and a maximum injection time of 50 ms. Spectra with unassigned charges, and charges of +1 and >8 were excluded from the precursor selection.

The four datasets (lysates of pRMG and MIO-M1 cells, and secretomes of pRMG and MIO-M1 cells) were analyzed separately. The Proteome Discoverer 2.4 SP1 software (version 2.4.1.15; Thermo Fisher Scientific) was used for peptide and protein identification via a database search (Sequest HT search engine) against the SwissProt Human (MIO-M1) and Ensembl Pig (pRMG). Database search was performed with full tryptic specificity, allowing for up to one missed tryptic cleavage site, a precursor mass tolerance of 10 ppm, and fragment mass tolerance of 0.02 Da. Carbamidomethylation of Cys was set as a static modification. Dynamic modifications included deamidation of Asn and Gln, oxidation of Met, and a combination of Met loss with acetylation on protein N-terminus. Peptide spectrum matches and peptides were validated with the Percolator algorithm (42). Each dataset was measured as triplicate. One control sample in the dataset of MIO-M1 secretomes had an unexpected low overall abundance and was excluded as an outlier. Only the top-scoring hits for each spectrum were accepted with a false discovery rate (FDR) <1% (high confidence). The final list of proteins satisfying the strict parsimony principle included only protein groups passing an additional protein confidence filter FDR <5% filter (target/decoy concatenated search validation).

Quantification of proteins, after precursor recalibration, was based on intensity values (at RT apex) for all unique peptides per protein. Peptide abundances were normalized to the total peptide amount. The protein abundances were calculated summing the normalised abundance values of corresponding unique peptides. These protein abundances were used for calculation of enrichment ratios of proteins in the stimulated samples to the untreated samples, resulting in single ratios for every quantified protein in every treated sample. Significance of the ratios was tested using a background-based t-test with correction for multiple testing according to Benjamini-Hochberg (adjusted *P*-value) (43, 44).

### 2.5 Data analysis and visualization

Calculation of the abundance ratio weight requires abundance values for both, the stimulated sample and the control. However, if a protein is exclusively expressed in one of these samples, the Proteome Discoverer software fails to calculate a respective abundance ratio weight. Since these extreme values were of special interest to us, the missing abundance ratio weights were imputed using the R package mice (version 3.13.0) and the “classification and regression trees” imputation method.

Ingenuity Pathway Analysis (IPA; Qiagen, Hilden, Germany) was used to analyze overrepresentation of proteins in canonical pathways of the IPA library, as described elsewhere (45). IPA allows deducing potential physiological effects of the various tested cytokines. Analysis was performed based on the fold-change of the stimulated samples and the abundance ratio *P*-value. Fisher’s exact test allowed testing for nonrandom associations of proteins in the datasets and the different canonical pathways (46). Furthermore, the method of Benjamini-Hochberg (B-H *P*-value) corrected for multiple testing (44).

The euclidean distance for the heatmap analysis was calculated with the open source software Cluster 3.0 and hierarchically clustered by complete linkage clustering (47). The resulting heat map was visualized with the open source software Java Treeview (version 1.2.0) (48).

## 3 Results

### 3.1 Differential secretion of proteins after stimulation of Müller cells with various cytokines

Müller cells are in close contact to all retinal cells, the vitreous and the blood vessels (17). To address, whether this privileged position within the retina also translates into extensive signaling between Müller cells and the surrounding cells, the secretomes of the human Müller cell-derived cell line MIO-M1 and of pRMG were quantitatively analyzed by mass spectrometry after stimulation for 24h with the cytokines IFNγ, IL-10, IL-4, IL-6, TGFβ1, TGFβ2, TGFβ3, TNFα and VEGF, respectively. By this means, we quantified 2,031 proteins in the supernatant of MIO-M1 cells (Suppl. Table 1) and 3,093 proteins in the supernatant of pRMG across all treatment groups (Suppl. Table 2).

Fig. 1 and Suppl. Fig. S1 summarize changes in the secretome after treatment of Müller cells with the different cytokines. A log_2_ fold change of ±0.58 and a corrected p-value of equal or less than 0.05 served as cutoff to define significantly upregulated or downregulated genes, respectively. Proteins equally regulated in MIO-M1 cells and pRMG were labeled with their gene symbol. However, this was only possible for proteins with identical gene symbols in the human and the porcine protein database. After treatment with IFNγ, 107 proteins in the secretome of MIO-M1 cells and 176 proteins in the secretome of pRMG were significantly more abundant, while 67 proteins of MIO-M1 cells and 96 proteins of pRMG were significantly less abundant in the supernatants (Fig. 1A; Suppl. Fig. S1A). Intriguingly, MIO-M1 cells and pRMG shared 21 upregulated and one downregulated protein. Among these shared regulated proteins after treatment with IFNγ were many with immune system functions, like signaling molecules (e.g. C-X-C Motif Chemokine Ligand 9 (CXCL9), CXCL10, IL-6) and components of the complement system (e.g. C1R, Serpin Family G Member 1 (SERPING1)). Upon treatment with TNFα, 127 (MIO-M1) or 143 (pRMG) proteins were more abundant and 57 (MIO-M1) or 87 (pRMG) proteins were less abundant in the supernatant (Fig. 1B, Suppl. Fig. S1B). Within these groups, MIO-M1 cells and pRMG shared 20 upregulated and three downregulated proteins, again with many pro-inflammatory proteins like C-X3-C Motif Chemokine Ligand 1 (CX3CL1), CXCL10, C-C Motif Chemokine Ligand 2 (CCL2), IL-6, and C1R being upregulated. Thus, IFNγ and TNFα resulted in the most conserved changes of the secretome of Müller cells when comparing between stimulated MIO-M1 cells and pRMG. In contrast, MIO-M1 cells and pRMG only shared ten proteins that were more abundant and three which were less abundant after treatment with VEGF, with seven proteins being keratins (Fig. 1C, Suppl. Fig. S1C). Treatment of Müller cells with the three different interleukins had a subtle influence on secreted proteins with conserved regulation of seven (IL-4), nine (IL-6), and two (IL-10) proteins in the secretome of MIO-M1 cells and pRMG (Fig. 1D-F, Suppl. Fig. S1D-F).

**Figure 1.**
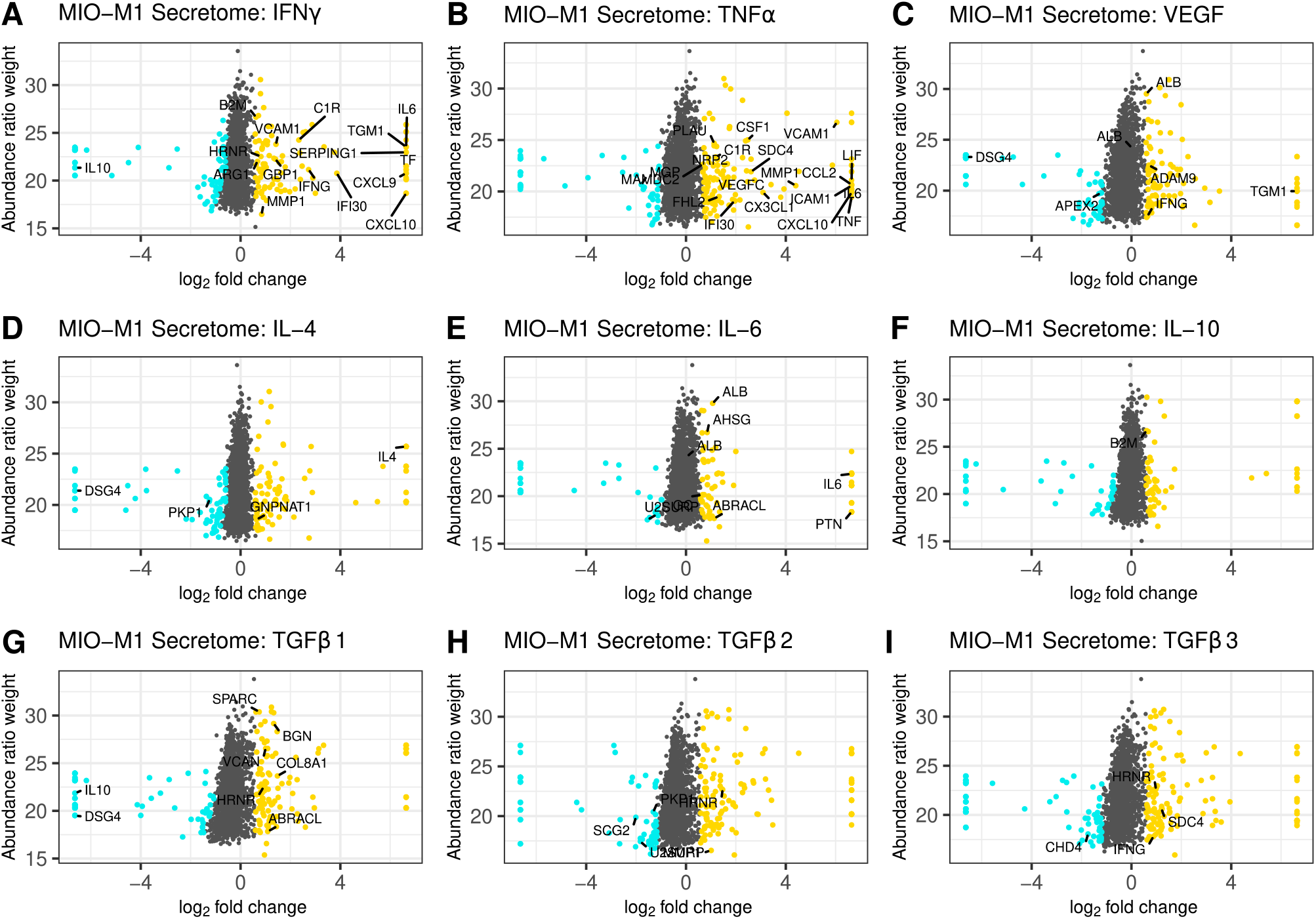
Scatterplot of all identified proteins from MIO-M1 secretomes after treatment with the indicated cytokines for 24 h (**A-I**). Proteins with significant changes in their abundance (±log_**2**_(1.5) fold expression, corrected p-value ≤0.05) were colored, with upregulated proteins being depicted as yellow dots, while down-regulated proteins are colored cyan. Proteins with significantly altered abundance in both, MIO-M1 and pRMG secretomes, are labeled with their gene symbol. Keratins were excluded.

Finally, TGFβs led to pronounced alterations in the secretome of MIO-M1 cells. TGFβ1 enhanced the secretion of 125 proteins while simultaneously reducing the abundance of 67 proteins, TGFβ2 increased the abundance of 131 proteins while impairing secretion of 69 proteins, and TGFβ3 raised abundance of 135 proteins and reduced the abundance of 76 proteins (Fig. 1G-I). Furthermore, eleven proteins of MIO-M1 cells and pRMG were similarly regulated by TGFβ1, eight upregulated and three downregulated (Suppl. Fig. S1G). After treatment with TGFβ2, three proteins were more abundant and four proteins less abundant in the secretome of both cell types (Suppl. Fig. S1H). Additionally, the secretome of TGFβ3 treated MIO-M1 cells and pRMG shared one downregulated protein and eight upregulated proteins, with most proteins like Osteonectin (Secreted Protein Acidic And Rich In Cysteine; SPARC), Matrix Metallopeptidase 1 (MMP1), Biglycan (BGN) and various keratins being functionally related to extracellular matrix organization (Suppl. Fig. S1I). Furthermore, stimulation with TGFβ3 evoked upregulation of pro-inflammatory cytokine IFNγ in both, the MIO-M1 cell line and pRMG. Overall, there are subtle, but intriguing differences in protein abundances after stimulation of Müller cells with the different TGFβ isoforms. Some of the proteins differentially induced by the TGFβ isoforms in MIO-M1 cells are pro-inflammatory cytokines, like IFNγ, TNFα, and CCL2 (Suppl. Table 1).

Next, we aimed to examine the physiological functions of proteins secreted by Müller cells. Therefore, we ordered secreted proteins in distinct clusters, grouped by similar expression patterns. From the identified proteins of the MIO-M1 supernatant, all cytosolic contaminants were removed, resulting in a list that exclusively contained extracellular proteins. Furthermore, only those proteins that showed a significant change of abundance (corrected p-value ≤0.05 and ± 1.5-fold abundance) in at least one treatment were selected for further analysis. As a result, a hierarchical heat map and an associated dendrogram consisting of 171 proteins was generated using the log_2_ abundance ratio of treated and untreated control cells (Suppl. Fig. S2). Furthermore, clusters were highlighted depending on their position on the dendrogram (Fig. 2). To avoid single protein clusters, the proteins Polymeric Immunoglobulin Receptor (PIGR) and Lacritin (LACRT) were assigned to Cluster F, despite being on different branches of the dendrogram.

**Figure 2.**
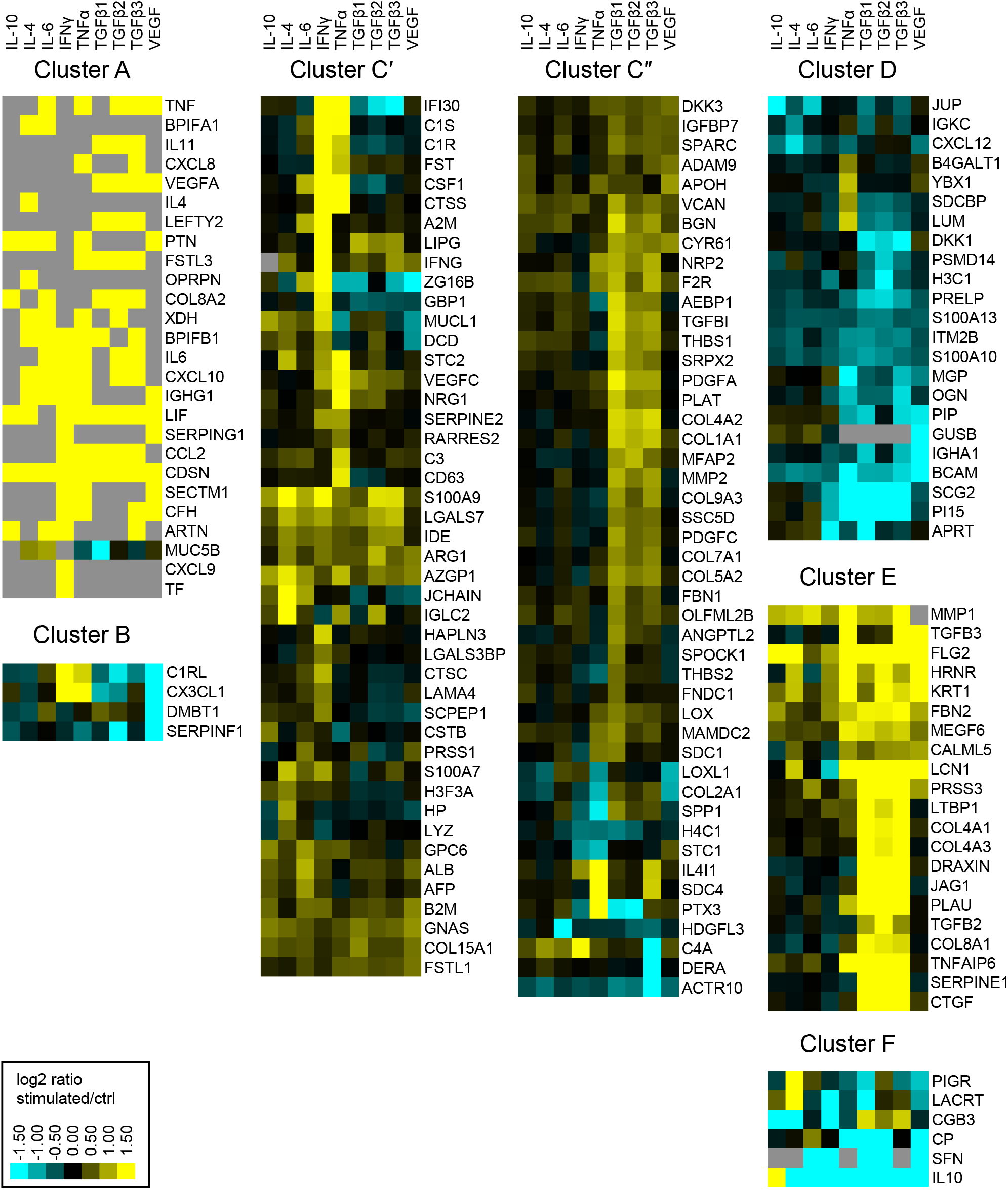
Heatmap of hierarchical cluster analysis of proteins secreted by MIO-M1 cells after treatment with various cytokines. Identified proteins were filtered for extracellular proteins with significant changes in expression (±log_**2**_(1.5) fold expression, corrected p-value ≤0.05). Down-regulated proteins are presented in cyan, while up-regulated proteins are depicted yellow for the respective treatments. Gray squares represent proteins that were neither identified in the untreated control, nor in the respective treatment. Clusters were defined using the branches of a dendrogram and shown as close up with the corresponding gene symbols.

Notably, we identified proteins which were secreted by MIO-M1 cells exclusively after stimulation (cluster A), while another set of proteins was completely absent in MIO-M1 cells (cluster B) treated with VEGF. The largest cluster comprises proteins upregulated by at least one cytokine (cluster C). In contrast to cluster D, consisting of proteins downregulated by TGFβs, TGFβs induce secretion of proteins of cluster E. Finally, secretion of most proteins of cluster F is deterred by at least one cytokine.

The clusters A, C and E are large clusters with proteins upregulated by at least one cytokine, with cluster A being the cluster with the most pronounced changes in secretion of its members and cluster C being the most diffuse one. In order to deduce the physiological functions of the secreted proteins of each of these clusters, we performed gene ontology (GO) analysis. Thereby, redundant pathways were condensed to a single representative pathway and the top 10 pathways with the lowest enrichment FDR were displayed. Intriguingly, the top 10 pathways for cluster A were related to immune system processes, with “Humoral immune response” and “Immune system process” being the most significant pathways with an enrichment FDR of 3.6 × 10^−9^ and 6.9 × 10^−9^, respectively (Fig. 3A). Furthermore, cluster C contained the pathway “Immune system process”, while most other pathways in this cluster C were involved in shaping the extracellular environment (Fig. 3B). Besides further pathways involved in extracellular remodeling, the proteins of cluster E were also associated to the pathway “AGE-RAGE signaling pathway in diabetic complications” (Fig. 3C).

**Figure 3.**
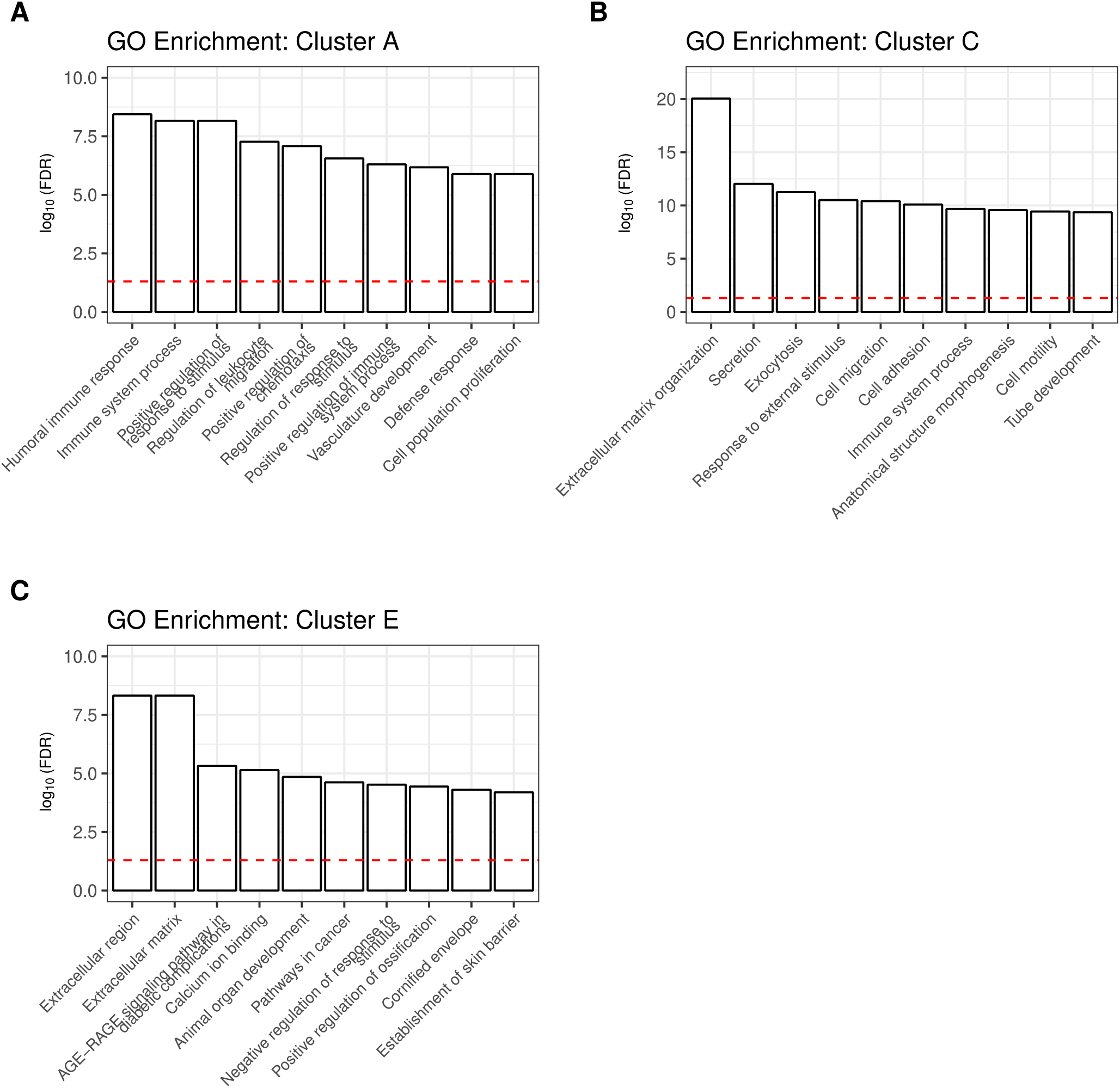
A shiny GO enrichment analysis for the previously defined cluster A (**A**), cluster C (**B**) and cluster E (**C**) was performed. Redundant pathways were reduced to show only one representative pathway. Depicted is the −log10(FDR) of the top 10 pathways for each cluster. The red dashed line indicates the significance threshold.

### 3.2 In-depth analysis of the Müller cell proteome after stimulation with a selection of cytokines

Our secretome analysis hints towards extensive signaling between Müller cells and their cellular environment, differentially induced by various cytokines. To elucidate the underlying cellular alterations, we also investigated differences in the proteome of MIO-M1 cells and pRMG cells by mass spectrometry after treatment with these cytokines for 24 h. In total, 5,514 proteins were quantified in the lysates of MIO-M1 cells (Suppl. Table 3) and 4,187 proteins in the lysates of pRMG cells (Suppl. Table 4) across all treatment groups.

The threshold for significant abundance changes was set using the same cutoff values as for the secretome, and equally regulated proteins in MIO-M1 cells and pRMG were labeled with their gene symbol, if they shared the same gene symbol in the human and the porcine database (Fig. 4; Suppl Fig. S3). Although the porcine protein database contains mostly humanized gene symbols, the swine leukocyte antigens (SLA) genes show little sequence homology between the human and the porcine genome and cannot properly be humanized (49). However, Human Leukocyte Antigen-C Alpha Chain (HLA-C) was part of the porcine protein database.

**Figure 4.**
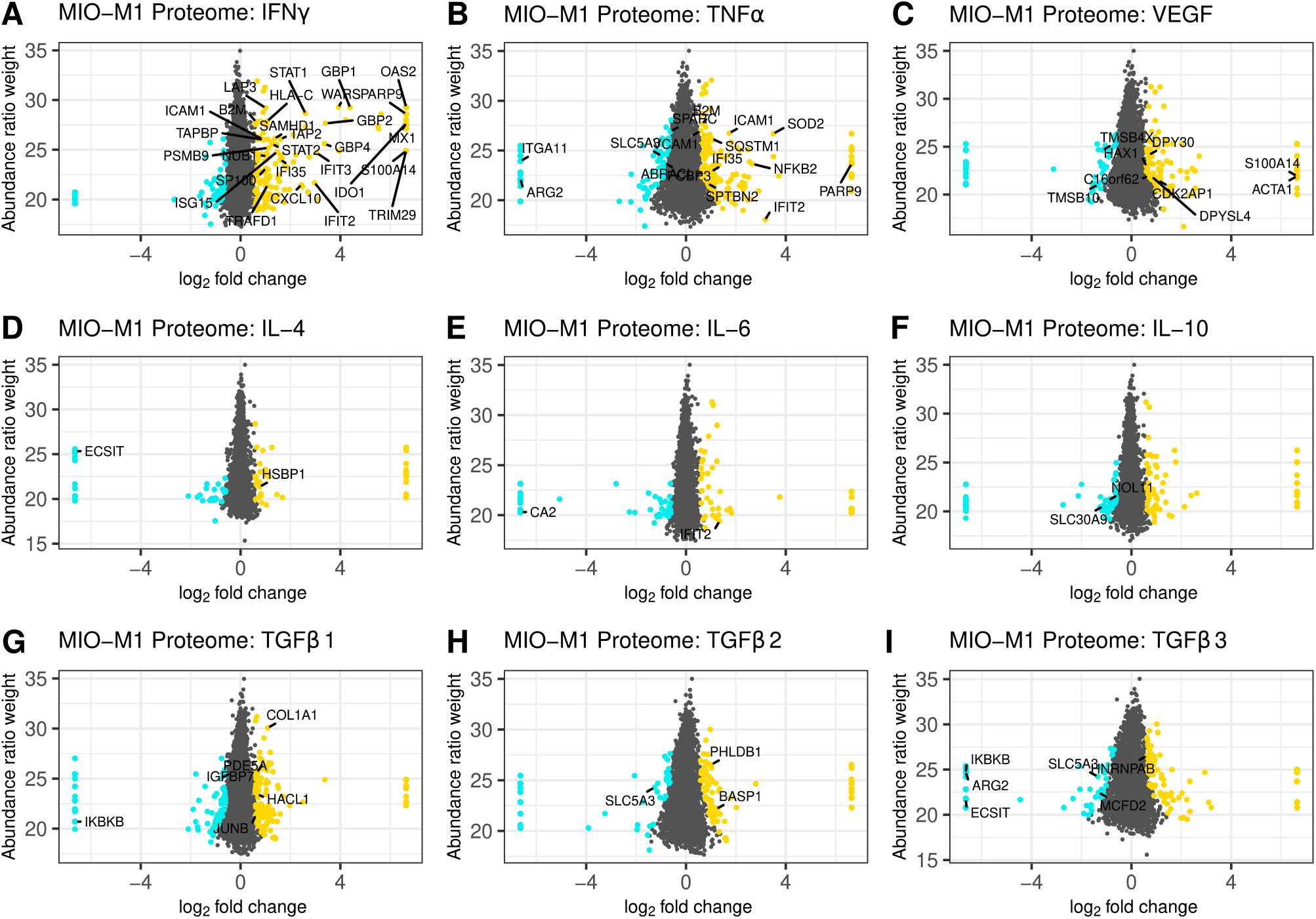
Scatterplot of all identified proteins from MIO-M1 lysates after treatment with the indicated cytokines for 24 h (**A-I**). Proteins with significant changes in their abundance (±log_**2**_(1.5) fold expression, corrected p-value ≤0.05) were colored, with upregulated proteins being depicted as yellow dots, while down-regulated proteins are colored cyan. Proteins with significantly altered abundance in both, MIO-M1 and pRMG lysates, are labeled with their gene symbol. Keratins were excluded.

Treatment of Müller cells with IFNγ resulted in 206 more abundant and 88 less abundant proteins in MIO-M1 cells and 331 more abundant and 36 less abundant proteins in pRMG lysates (Fig. 4A; Suppl Fig. S3A). Thereof, 29 proteins showed higher expression levels in both cells types. Among the overlapping proteins were restriction factors like SAM And HD Domain Containing Deoxynucleoside Triphosphate Triphosphohydrolase 1 (SAMHD1) and MX Dynamin Like GTPase 1 (MX1), transcription factors like Signal Transducer And Activator Of Transcription 1 (STAT1) and STAT2, regulators of protein homeostasis like Leucine Aminopeptidase 3 (LAP3) or the Proteasome 20S Subunit Beta 9 (PSMB9), and proteins associated with peptide transport and antigen presentation like Transporter 2, ATP Binding Cassette Subfamily B Member (TAP2), TAP Binding Protein (TAPBP), Beta-2-Microglobulin (B2M), as well as HLA-C. After treatment with TNFα, 204 proteins were more abundant in the proteome of MIO-M1 cells, while 119 proteins were less abundant (Fig. 4B). In pRMG, 207 proteins with higher abundance and 285 proteins with lower abundance were identified upon treatment with TNFα, with 18 proteins that were differentially regulated in both cell types (Suppl. Fig. S3B). Among shared proteins that were more abundant after treatment with TNFα were pro-inflammatory proteins like B2M and Nuclear Factor Kappa B Subunit 2 (NFKB2), or adhesion molecules like Intercellular Adhesion Molecule 1 (ICAM1) or Vascular Cell Adhesion Molecule 1 (VCAM1). VEGF led to 143 more and 102 less abundant proteins in MIO-M1 cells or 232 more and 224 less abundant proteins in pRMG, respectively (Fig. 4C; Suppl. Fig. S3C). Thereof, MIO-M1 cells and pRMG shared nine more abundant proteins, inter alia proteins associated with reorganization of the cortical cytoskeleton like Alpha-Actin-1 (ACTA1) or HCLS1 Associated Protein X-1 (HAX1), and two less abundant proteins, Thymosin Beta 10 (TMSB10) and Thymosin Beta 4 X-Linked (TMSB4X), both inhibitors of actin polymerization. Upon treatment with interleukins IL-4, IL-6 and IL-10, the Müller cell proteomes mirrored the subtle effects of these cytokines on the abundance of proteins observed for the Müller cell secretomes (Fig. 4D-F, Suppl. Fig. S3D-F). Also in line with the secretome data, the overlap between differentially abundant proteins of the MIO-M1 and pRMG proteome after treatment with the various interleukins contained only few proteins. In contrast, TGFβ1 increased the abundance of 143 proteins, while decreasing the abundance of 94 proteins in the proteome of MIO-M1 cells and increased the abundance of 203 proteins, while decreasing the abundance of 103 proteins in the proteome of pRMG (Fig. 4G; Suppl. Fig. S3G). In comparison to the lower abundant proteins Phosphodiesterase 5A (PDE5A) and Inhibitor Of Nuclear Factor Kappa B Kinase Subunit Beta (IKBKB), the proteins Collagen Type I Alpha 1 Chain (COL1A1), Insulin Like Growth Factor Binding Protein 7 (IGFBP7), JunB Proto-Oncogene (JUNB), and 2-Hydroxyacyl-CoA Lyase 1 (HACL1) were more abundant in both, MIO-M1 cells and pRMG after treatment with TGFβ1. Following treatment with TGFβ2, 125 proteins of the proteome of MIO-M1 cells and 266 proteins of the proteome of pRMG were more abundant, whereas 67 proteins of the MIO-M1 proteome and 229 proteins of the pRMG proteome were less abundantly expressed (Fig. 4H; Suppl. Fig. S3H). In the case of treatment with TGFβ3, 130 proteins in the MIO-M1 proteome and 185 in the pRMG proteome showed higher abundances, while 94 proteins in MIO-M1 proteome and 250 in the pRMG proteome were less abundant (Fig. 4I; Suppl. Fig. S3I). The overlap of MIO-M1 cells and pRMG treated with TGFβ2 comprised three proteins, and treatment with TGFβ3 resulted in an overlap of seven proteins. Overall, pRMG reacted more pronounced to treatment with the various cytokines compared to MIO-M1 cells.

### 3.3 Canonical pathways enriched in Müller cells upon stimulation

Treatment with various cytokines partly induced pronounced changes in the secretome and proteome of Müller cells. In the secretome, these changes primarily included the secretion of pro-inflammatory cytokines and proteins associated with organization of the extracellular matrix. To elucidate overrepresented mechanisms and pathways in stimulated Müller cells, we performed Ingenuity pathway analysis (IPA). We limited the IPA to significantly regulated proteins (p-value ≤ 0.05) identified in the MIO-M1 and pRMG lysates. Since IPA cannot handle porcine gene symbols, we replaced the only canonical SLA gene SLA-1 in our pRMG dataset by the canonical human HLA gene HLA-A. Hence, an IPA core analysis was performed with 1,543 proteins for pRMG and with 2,262 proteins for the MIO-M1 cells. IPA identified 338 canonical pathways within the proteome of MIO-M1 cells and 218 canonical pathways in the proteome of pRMG that were significantly enriched by at least one of the used cytokines (IPA p-value ≤ 0.05; Suppl. Table 5). Among the identified canonical pathways were many pathways associated with signaling, cell death, immune system processes and the cellular redox state (Fig. 5; Suppl. Fig. S4). A selection of canonical pathways enriched in both pRMG and MIO-M1 cells after treatment with at least one of the tested cytokines is depicted in Fig. 5.

**Figure 5.**
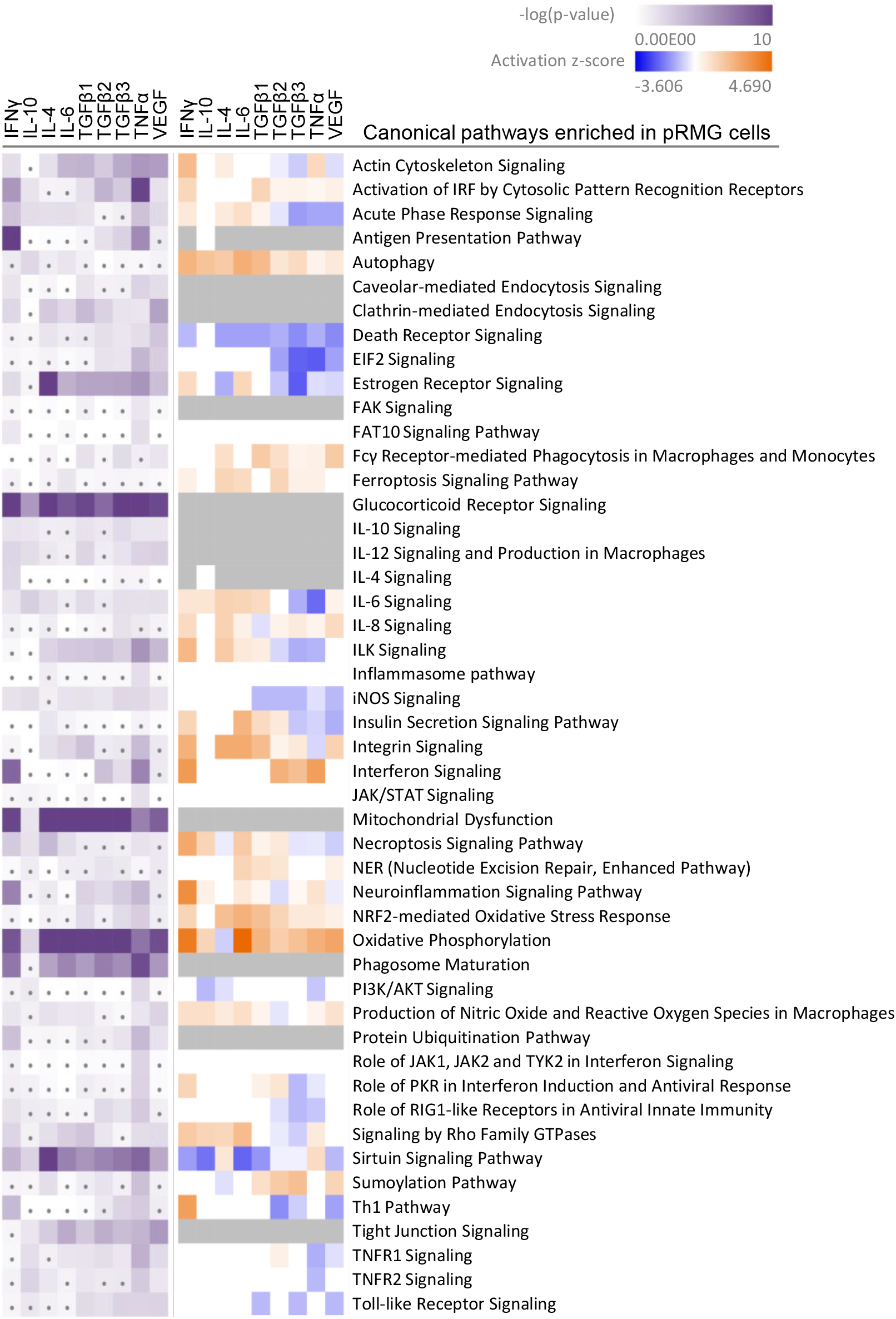
A comparative IPA analysis with the significantly regulated proteins identified in the pRMG lysates after stimulation with the indicated cytokines was performed. Canonical pathways related to signaling, cell death, immune system processes and oxidative stress were selected. Pathways with significant enrichment of genes after stimulation with at least one cytokine are presented. Significance of the gene enrichment for each pathway and treatment is indicated by purple squares in the left array. Thereby, treatments that did not meet the significance threshold (p-value ≤ 0.05) are marked with a dot. The z-score is indicated in the right array and represents a prediction of activation (orange) or inhibition (blue) of the pathway. Gray squares mark treatments where the activation state of a pathway could not be calculated.

Among the most significantly regulated pathways in MIO-M1 cells and pRMG were the canonical pathways “Mitochondrial Dysfunction” and “Oxidative Phosphorylation”. These pathways were significantly induced by all examined cytokines in pRMG. The pathways “Ferroptosis Signaling Pathway”, “iNOS Signaling”, “NRF2-mediated Oxidative Stress Response”, and “Production of Nitric Oxide and Reactive Oxygen Species in Macrophages” are closely linked to the cellular redox state and were among the enriched pathways in MIO-M1 cells and pRMG after treatment with various cytokines. Furthermore, proteins associated with the maturation of phagosomes were significantly enriched in pRMG. In line with this, “Caveolar-mediated Endocytosis Signaling” was significantly enriched in pRMG after treatment with IFNγ, TGFβ1, TNFα and VEGF, and “Clathrin-mediated Endocytosis Signaling” was significantly enriched in pRMG after treatment with all cytokines except IL-10. Besides these two pathways associated with the recycling of the extracellular environment, intracellular protein homeostasis and MHC class I peptide generation was facilitated by enrichment of the “Protein Ubiquitination Pathway” in pRMG after treatment with IFNγ, TGFβ3, TNFα and VEGF. Similarly, IFNγ, IL-4, TGFβ1, TGFβ3, TNFα and VEGF enriched “Protein Ubiquitination Signaling” in MIO-M1 cells. “Neuroinflammation Signaling” was induced by IFNγ, TNFα and VEGF in MIO-M1 cells and by IFNγ, TGFβ1, TGFβ3 and TNFα in pRMG, whereas TGFβ2 and VEGF led to a slight inhibition of this pathway in pRMG.

### 3.4 A deeper look into Müller cell complement secretion upon cytokine stimulation

Because the enrichment analysis of the secretome yielded highly significant hits such as “humoral immune response” and “immune system process,” we took a closer look at complement proteins in the secretome and cell lysates. Notably, most complement proteins are secreted as key components of the humoral immune system. The identified complement components include central complement proteins, regulators, and receptors. Consistent with their localization in the cell membrane, the latter (including ITGAM, ITGB2, C5aR1) were detected only in cell lysates, and here specifically in those of pRMGs. The complement regulators clusterin (CLU), vitronectin (VTN), CD59, and SERPING were found in most test samples. With regard to the central complement components, the pRMG secretome took a prominent position and showed results for complement components for all three different activation pathways (e.g. C1q, FD, MASP1) and the terminal pathway (e.g. C9). The central complement protein C3 was found in both the MIO-M1 and pRMG secretomes and in the MIO-M1 lysate. Interestingly, cytokine treatment induced changes in complement proteins and regulators but had no effect on complement receptor expression. We observed that C1q subunits, which initiate the classical complement pathway by binding to antibodies, were detectable only in pRMG but not in MIO-M1 cells. C1q levels in cell lysates and the corresponding secretome were consistently reduced after TNFα treatment but were increased by IFNγ. Moreover, complement proteases C1r and C1s, which bind to C1q therewith continuing the cascade of classical pathway activation, were enriched in the supernatants of MIO-M1 and pRMG cells treated with IFNγ (Fig. 2C). In contrast, C1r concentration was significantly decrease in supernatants of MIO-M1 cells but not pRMG after VEGF and TGFβ2 application. Notably, C1s and C1r were not detected in cellular lysates. Interestingly, the abundance of the central complement proteins C3 and C4A were modified by the supplemented cytokines in MIO-M1 secretomes only and not in any other data set (Fig. 2C). These proteins are cleaved upon complement activation as for example triggered by the C1q-mediated classical pathway and result in cleaved products which interact with cellular receptors (e.g. C3a/C3b, C4a). Here, complement protein C3 is mainly increased following TNFα addition and C4 upon exposure to IFNγ (Fig. 2C). In fact, IFNγ was also the major player modulating the secreted complement components in pRMG: C2 and FI were significantly increased while C9, FD and MASP1 were clearly reduced in its presence. These complement components absent from any other sample.

Regarding the complement regulators factor H (FH), SERPING and CLU are of interest. Secretion of FH was not observed in untreated MIO-M1 and pRMG, but it was significantly upregulated in MIO-M1 secretomes following IFNγ, TNFα, TGFβ3, and VEGF treatment (Fig. 2A). Similar results were obtained for SERPING, whose levels were increased by IFNγ in the MIO-M1 secretome, pRMG cell lysates and secretome. Remarkably, the MIO-M1 lysate showed decreased values for CLU following IFNγ, TGFβ1 and TNFα, and similar but not significant trends was observed for the respective secretome. Finally, while CLU was upregulated in pRMGs lysates upon IL-6 or VEGF treatment, no significant alterations could be found in corresponding secretomes.

In summary, IFNγ and TNFα seemed to be the most effective cytokines to modulate the Müller cell complement expression and secretion (Fig. 2).

### 3.5 Müller cells as atypcial antigen-presenting cells

Intriguingly, treatment of pRMG with IFNγ, TGFβ2, TGFβ3 and TNFα significantly enriched proteins associated with the “Antigen Presentation Pathway”. Likewise, the “Antigen Presentation Pathway” was induced in MIO-M1 cells by treatment with IFNγ, TGFβ1, TNFα and VEGF. Thereby, antigen presentation is an umbrella term for two distinct processes. MHC class I antigen presentation is common to all nucleated cells and allows CD8^+^cytotoxic T cells (CTL) to assess whether cells are infected with an intracellular pathogen (50, 51). In contrast, MHC class II is presented to antigen specific CD4^+^T cells mainly by professional antigen-presenting cells inducing their activation and differentiation to T helper cells (52).

To investigate the antigen presentation capacity of Müller cells, we constructed a hierarchical heatmap for MIO-M1 cells (Fig. 6A) and pRMG (Fig. 6B) challenged with various cytokines. Proteins linked to antigen presentation were selected and clustered hierarchically. Proteins associated with MHC class I antigen presentation are displayed in the upper panel and proteins correlated to MHC class II antigen presentation are depicted in the lower panel. In MIO-M1 cells, IFNγ induced the majority of proteins linked to both, MHC class I and II antigen presentation, whereas TNFα exerted its inductive effect exclusively on MHC class I antigen presentation. The only protein linked to MHC class II antigen presentation induced by TNFα in MIO-M1 cells was Cathepsin S (CTSS). Furthermore, the other cytokines did not induce proteins related to antigen presentation in MIO-M1 cells. Quite the contrary, TGFβ2 and TGFβ3 reduced the abundance of proteins linked to MHC class I antigen presentation in these cells. In contrast to MIO-M1 cells, pRMG reacted to all tested cytokines by induction of components of both MHC class I and II antigen presentation to varying degrees. IFNγ and TNFα induced proteins of MHC class I and II antigen presentation in pRMG, among others SLA-DQA1, SLA-DQB1, SLA-DRA and SLA-DRB1. Furthermore, class I and II antigen presentation was upregulated by TGFβ isoforms 1-3 in pRMG. While TGFβ2 and TGFβ3 induced the components of the MHC class I peptide loading complex TAP2 and TAPBP, TGFβ1 also increased the abundance of HLA-DMA and HLA-DMB, proteins involved in the peptide loading on MHC class II. We saw only a subtle induction of proteins related to antigen presentation by IL-10, IL-4 and IL-6. The smallest impact on antigen presentation proteins was seen after stimulation with IL-6. Furthermore, IFNγ significantly upregulated the expression of the co-stimulatory molecule CD40 in pRMG, while TGFβ2, TGFβ3, TNFα and VEGF resulted in lower abundance of CD40 (Suppl. Table 4).

**Figure 6.**
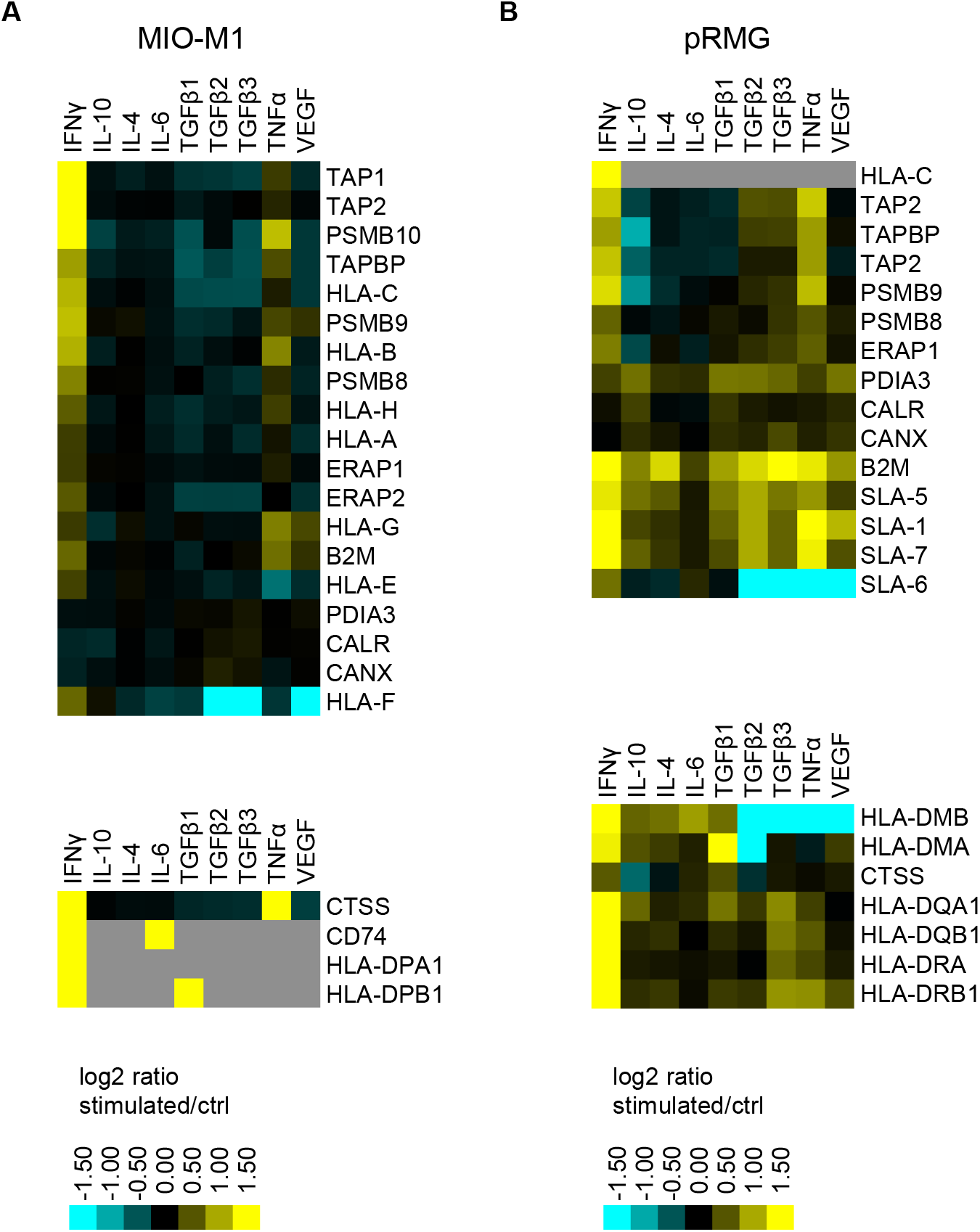
Heatmap of hierarchical cluster analysis of proteins involved in MHC class I (upper panel) and MHC class II (lower panel) antigen presentation expressed by MIO-M1 cells (**A**) and pRMG cells (**B**) after treatment with various cytokines. Down-regulated proteins are presented in cyan, while up-regulated proteins are depicted yellow for the respective treatments. Gray squares represent proteins that were neither identified in the untreated control, nor in the respective treatment. The heatmap was generated on the basis of the log_**2**_ fold change of the respective proteins.

Induction of the canonical MHC class I and MHC class II antigen presentation pathway as assessed by IPA for pRMG after treatment with IFNγ is summarized in Fig. 7. This pathway was enriched in pRMG cells with a p-value of 4.15 × 10^−12^. Besides MHC class I and MHC class II, also components of the peptide loading complex of MHC class I (TAP2 and TPN) were upregulated in pRMG after IFNγ treatment. Furthermore, IFNγ induces the Large Multifunctional Peptidase 2 (LMP2; synonymous to PSMB9) and Large Multifunctional Peptidase 7 (LMP7; synonymous to PSMB8) subunits of the immunoproteasome in MIO-M1 cells.

**Figure 7.**
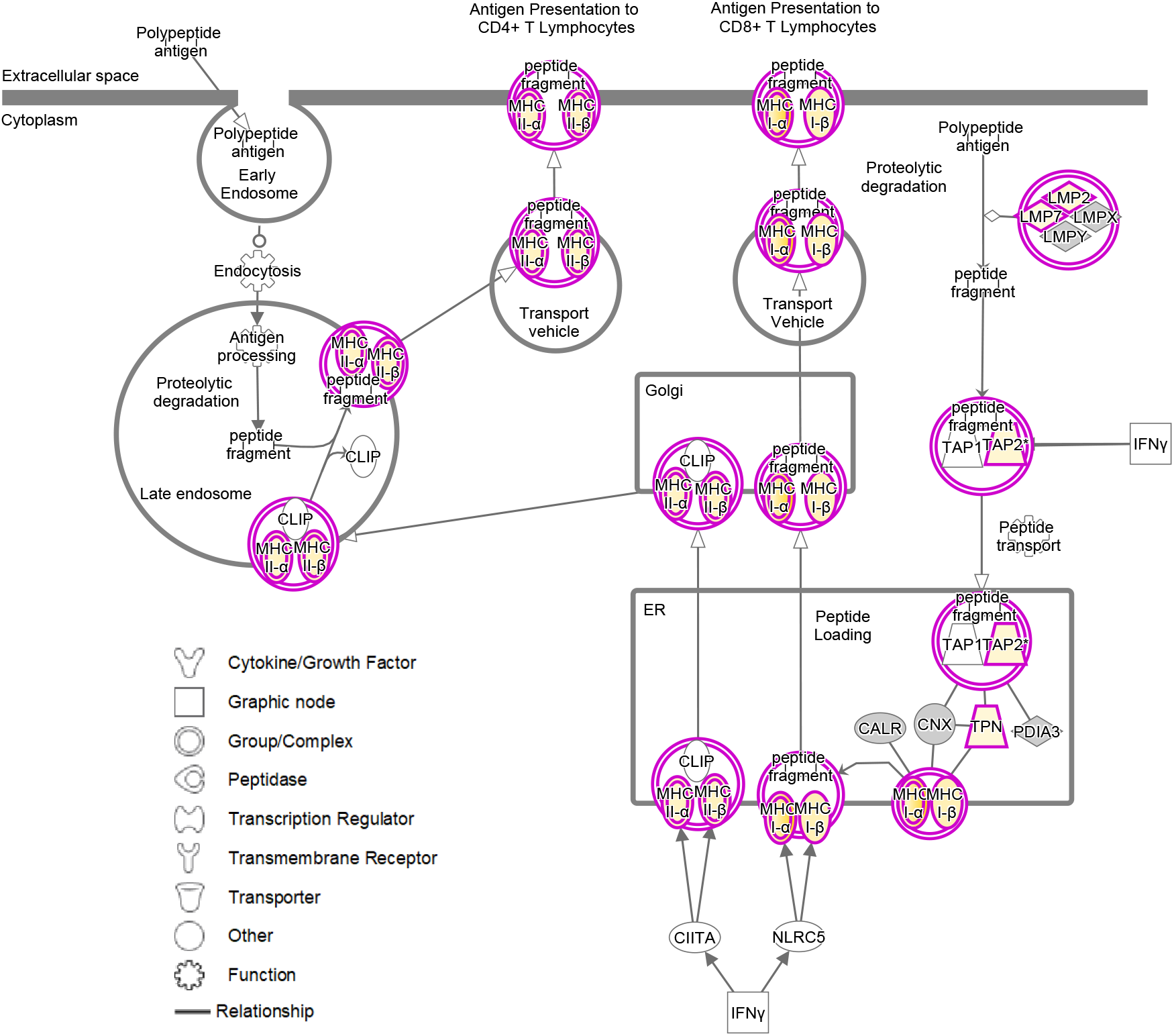
IPA for pRMG cells after treatment with IFNγ was performed. Depicted is the canonical antigen presentation pathway. Yellow proteins are induced. The intensity of the yellow color indicates the degree of upregulation. Grey proteins are in the dataset but did not pass the analysis cutoffs of the pathway analysis. Purple circles highlight proteins (single circle) or complexes (double circles).

## 4 Discussion

DR is a condition associated with microvascular degeneration, resulting in ocular inflammation and eventually in complete blindness (53). As the prevalence of diabetes in developed and developing countries rises, DR became the leading cause of blindness in the working-age population (54). Microglia are resident innate immune cells in the retina and as such, they are considered to regulate the inflammation during DR (55). In contrast, the role of Müller cells during the pathology of DR has been often described to be the metabolic support of neurons (53). However, growing evidence suggests excessive signaling between Müller cells and the surrounding tissue including microglia, with both beneficial and detrimental consequences for the pathogenesis of DR (56). We show here that stimulation of Müller cells with pro-inflammatory cytokines like IFNγ or TNFα, but also with growth factors like TGFβ or VEGF resulted in profound, yet distinct response profiles in their secretomes, thus confirming their central role in cell-to-cell communication within the retina.

Previously, it has been shown that the conditional knock-out of VEGF in Müller cells of mice reduced the expression of inflammatory markers like TNFα or ICAM1 compared to diabetic control mice, indicating a pro-inflammatory effect of VEGF on Müller cells (57, 58). Here we could show that VEGF induced the Müller cell line MIO-M1 to secrete proteins associated with immune effector processes like IFNγ, B2M, HRNR, and complement-associated proteins like SERPING1 and complement factor H (FH). Furthermore, ICAM1 was more abundant in our proteome data of MIO-M1 cells and pRMG treated with VEGF compared to the respective control cells. Thus, we were able to confirm the pro-inflammatory role of VEGF in Müller cells.

Overall, we observed that MIO-M1 cells secreted TNFα upon stimulation with VEGF, IL-6, and the TGFβ isoforms 2 and 3. Increased levels of TNFα were previously identified in the vitreous of DR patients (59). Furthermore, Müller cells have been reported to secrete increased amounts of VEGF and TNFα under stress conditions like inflammation or hyperglycemia, thereby promoting retinal inflammation (57, 60–64). Our results show that TNFα induced MIO-M1 cells to secrete inflammatory proteins but also proteins associated with tissue development like Leukemia Inhibitory Factor (LIF), Follistation (FST), Neuregulin 1 (NRG1), or Pleiotrophin (PTN), to name but a few. Thus, our data suggest a pleotropic role of TNFα in Müller cells.

TGFβs regulate early embryogenesis, maintenance and regeneration of mature tissues, and various disease processes (65). TGFβ1 has been described to be anti-inflammatory, inducing immune tolerance in the periphery and preventing autoimmunity. Thus, knock out of TGFβ1 in mice induced autoantibodies and multifocal inflammatory disease in many tissues, eventually causing the mice to die within 4 weeks of age (66). In our *in vitro* model, treatment of MIO-M1 cells with TGFβ1 induced secretion of CCL2 and therefore might result in the recruitment of leukocytes.

Simultaneously, TGFβ1 enhanced the secretion of anti-inflammatory proteins like TNF Alpha Induced Protein 6 (TNFAIP6) or LIF in Müller cells. In contrast, TGFβ2 and TGFβ3 also promoted immune system processes in MIO-M1 cells by inducing secretion of pro-inflammatory cytokines TNFα, CXCL8, CXCL10 and IFNγ. It has been shown that TGFβ2 acts immunomodulatory in aqueous humor by decreasing the expression of IL-6, CXCL1, CCL2, G-CSF and IGFBP-5 (67). However, we demonstrated that treatment of Müller cells with TGFβ2 induced expression of IL-6 and CCL2, suggesting a complex and diverse role of TGFβ2 signaling in the eye. Although TGFβ isoforms are closely related (sharing 71-79% sequence identity), have similar three-dimensional structures and signal canonically through the same receptor (68), our secretome analyses provide evidence that they affect Müller cells differentially. While knock-out of the TGFβ isoforms resulted in different phenotypes, all null mutations were lethal (69–72). Replacement of the coding sequences by one of the other isoforms only partially rescued these phenotypes, implying intrinsic differences of these isoforms (73, 74). These differences might originate from varying flexibilities of an interfacial helix and thus modify the binding of the TGFβ isoforms to matrix proteins as suggested by Huang and colleagues (68). Latent TGFβ binds Thrombospondin 1 (THBS1), augmenting its activation (75–77). Intriguingly, in our study, all three TGFβs elevate the abundance of THBS1 indicating a positive feedback loop. Recently, transcriptome analysis linked the expression of TGFβ isoforms of mice to activation of different signaling cascades. While TGFβ1 and TGFβ2 evoked the non-canonical p38MAPK signaling pathway, which has been linked to gliosis, TGFβ3 induced the SMAD canonical signaling pathway for TGFβs in mice (78, 79). During glaucoma, the aqueous humor contains elevated levels of TGFβ2, exceeding levels for homeostatic signaling (77, 80). Thereby, TGFβ2 has been associated with pathological remodeling of the trabecular meshwork and the optical nerve head (77). Furthermore, it stimulated secretion of extracellular matrix proteins by astrocytes and cells of the lamina cribrosa (81–84). Our data suggest that Müller cells also contribute to the remodeling of the extracellular matrix as stimulation with TGFβ1 and TGFβ3 resulted in enhanced secretion of extracellular matrix proteins like Fibrillin-2 (FBN2), various keratins, and collagens. Furthermore, they simultaneously enhanced the turnover of extracellular matrix by inducing the secretion of the Matrix Metalloprotease-1 (MMP1) and MMP2. Previously, MMP2 has been described to be elevated in DR patients and to promote the pathogenesis of DR by inducing mitochondrial dysfunction and apoptosis of retinal capillary cells (85). Intriguingly, our IPA also indicates that the canonical pathway for mitochondrial dysfunction is enriched in Müller cells upon stimulation with all TGFβ isoforms. However, this is not solely due to the enhanced secretion of MMP2, since mitochondrial dysfunction is enriched in Müller cells by all tested cytokines, whereas TGFβs exclusively induce MMP2.

A meta-analysis of eleven studies recently showed significantly increased levels of the anti-inflammatory cytokine IL-10 in the vitreous humor of DR patients (86). Moreover, IL-10 levels were higher in patients with proliferative DR compared to patients with non-proliferative DR (86). While our data show no contribution of Müller cells to elevated IL-10 secretion upon stimulation with the tested cytokines, IL-10 induced proteins associated with tissue development, cell adhesion, and angiogenesis in MIO-M1 cells, namely Corneodesmosin (CDSN), LIF, PTN, and various collagens, to name a few. In line with this, it has been shown that IL-10 is involved in the pathological angiogenesis during the development by modulating the macrophage response to hypoxia (87). Thus, we hypothesize that IL-10 might also be involved in the abnormal angiogenesis during DR.

Previously, it has been shown that the canonically anti-inflammatory IL-4 can potentiate cytokine and chemokine production in macrophages following a pro-inflammatory stimulus (88–90). In line with this, we observed increased abundances of many pro-inflammatory proteins like IFNγ, S100A9, S100A7, CXCL10, and Lysozyme C (LYZ) upon stimulation of MIO-M1 cells with IL-4, indicating that IL-4 does not exert an anti-but a pro-inflammatory influence in these cells. Interestingly, MIO-M1 cells did not secrete the anti-inflammatory interleukin IL-4 upon treatment with the various tested cytokines. Secretion of pro-inflammatory IL-6 by Müller cells has been described upon treatment with IL-1β or LPS (91). Furthermore, elevated levels of IL-6 were found in the vitreous and serum of DR patients and even further increased in patients suffering from proliferative DR (92). Here we show that IFNγ, TNFα, TGFβ2, and TGFβ3 also induced the secretion of IL-6 in MIO-M1 cells. Previously, it was shown that IL1β induced IL-6 through activation of the p38MAPK signaling pathway (93). The murine TGFβ isoforms 1 and 2 also activated the p38MAPK signaling in Müller cells of mice (78). Thus, the involvement of p38MAPK signaling in secretion of IL-6 by Müller cells should be addressed in further studies. Brandon and colleagues demonstrated induction of VEGF by IL-6 in Müller cells, especially under hyperglycemic conditions, preventing Müller cells from glucose toxicity (94). In contrast, our analysis revealed no VEGF secretion by MIO-M1 cells upon IL-6 treatment. High glucose concentrations of 25 mM potentiated the induction of VEGF by IL-6 (94). However, by default the standard culture medium used in our study has a D-glucose concentration of 25 mM. Thus, an adaption of MIO-M1 cells to these conditions might have occurred negating the induction of VEGF by IL-6.

Elevated levels of IFNγ in the vitreous of DR patients have been described previously (95, 96). In our study, we could observe distinctly elevated INFγ secretion only after stimulation with INFγ. Treatment with IFNγ furthermore induced the secretion of CXCL9, CXCL10, IL-6 and complement subcomponent C1R in MIO-M1 cells and pRMG.

Interestingly, we observed an induction of CX3CL1 in the whole-cell lysates and in the secretomes. Specifically, except for TGFβ1 and TGFβ2, all stimulants used in this study induced expression of CX3CL1 in the cell proteome of MIO-M1 cells, while only IFNγ and TNFα treatment resulted in significantly higher abundance of CX3CL1 in the secretome as well. CX3CL1 is a membrane-bound chemokine, which functions as an adhesion molecule for leukocytes, but can also be proteolytically cleaved, resulting in a soluble form with chemotactic function (97, 98). Several studies already demonstrated the implication of CX3CL1 in DR pathogenesis as the vitreous of PDR patients contains elevated levels of CX3CL1 (99) Furthermore, circulating CD11b+ leukocytes that are involved in leukostasis in DR express higher levels of CX3CR1 in diabetic mice compared to controls (100). Thus, membrane-bound CX3CL1 on the surface of Müller cells might be involved in leukostasis in DR. Since TNFα levels are elevated in the eyes of DR patients, this might result in the upregulation of CX3CL1 in diabetic Müller cells (101). In a previous study, incubation of microglia with Müller cell supernatant containing secreted CX3CL1 resulted in the upregulation of the respective receptor CX3CR1 in microglia. The authors proposed that Müller cells might be able to promote microglial motility via the chemotactic effect of CX3CL1 (102). Thus, secretion of CX3CL1 from Müller cells might contribute to DR pathogenesis by recruiting peripheral inflammatory cells and microglia. Upregulation of CX3CL1 after treatment with TNFα has been demonstrated in aortic endothelial cells, but could not be observed in retinal endothelial cells (103). Here, we demonstrate that stimulation with TNFα elicits secretion of CX3CL1 in pRMG and MIO-M1 cells and expression of CX3CL1 in MIO-M1 cells. Thus, we lay the ground for further research concerning the expression and regulation of CX3CL1 in Müller cells during DR.

Another route of communication between microglia/macrophages and Müller cells could be local changes in complement expression. We recently demonstrated, that Müller cells in the mouse retina are the main producers of complement components of the classical (C1s, C4), the alternative pathway (FB) and C3 as the central component to all pathways under homeostatic, but also under ischemic stress conditions (104). In the retina, it is the microglia that by far express highest levels of complement receptors including ITGAM (alias CD11b), C3aR, C5aR1 and C5aR2 (104). In our present study, TNFα and INFγ triggerd the most prominent effects on complement expression consistently in MIO-M1 and pRMG. Given that *in vivo* microglia and potentially other immune cell serve as major source of TNFα and INFγ (105) (resource data from Cowan et al., 2020 (106) at https://data.iob.ch), the strong effect of these cytokines on Müller cell could be central to coordinate the tissue immune homeostasis in pathologies. In this context, the enhanced expression of activating complement components by Müller glia could serve as feedback mechanisms towards microglia in turn modulating their activation profile. Taken together, our results point towards a pro-inflammatory phenotype of Müller cells, which is in line with a previous study, where we analyzed the surfaceome of primary equine Müller cells and MIO-M1 cells after stimulation with Lipopolysaccharide (30). While the surfaceome of Müller cells in this previous study revealed expression of MHC class I and II as well as costimulatory molecules especially in primary equine Müller cells, we could now further complement these results with LC-MS/MS-analyses of whole cell lysates and of cell supernatants, confirming an antigen-presenting phenotype of Müller cells. While stimulation with LPS resulted in enhanced expression of MHC class II molecules in primary equine Müller cells, no MHC class II molecules could be identified in MIO-M1 cells upon LPS treatment (30). In contrast, stimulation with IFNγ in this study induced the expression of proteins that are associated with both MHC-I and MHC-II antigen presentation in pRMG as well as in MIO-M1 cells. This is in accordance with an early study that demonstrated the induction of MHC class I and MHC class II molecules in primary human Müller cells by IFNγ *in vitro* (107). In contrast to MIO-M1 cells, pRMG also showed basal expression of MHC class II without stimulation. Thus, our data demonstrate that Müller cells exhibit several criteria for atypical antigen-presenting cells (108). While microglial cells are the predominant immune cells of the retina, our data confirm that Müller cells are crucially involved in immunological processes in the retina as well, as they possess an antigen processing and presenting machinery and secrete pro-inflammatory cytokines (14). We have previously shown that the cultivation of primary porcine Müller cells under hyperglycemic conditions resulted in higher expression levels of MHC class II molecules, pointing towards an immunologically activated state of Müller cells in DR (36).

Besides expression of MHC class I and II molecules, we demonstrated that stimulation of Müller cells with various cytokines resulted in the enrichment of proteins and pathways that are associated with the formation and maturation of phagosomes. Previously, Müller cells have been described to be phagocytic cells, capable of phagocytosing cell debris, dead photoreceptor cells and even bacteria (109–111). Our IPA showed that proteins of phagocytosis pathways in Müller cells are induced upon stimulation with various cytokines. Furthermore, phagocytosis is not only clathrin-but also caveolar-mediated. Since our data showed enrichment of phagocytic pathways, as well as the canonical antigen presentation pathway, it is possible that Müller cells present exogenous peptides on MHC class II to CD4+ T helper cells. Intriguingly, phagocytosis of dead photoreceptors would also allow Müller cells to present proteins expressed by photoreceptors on MHC class II, and via cross-presentation on MHC class I (112, 113). The relationship of HLA alleles and haplotypes to the progression of diabetic retinopathy is part of an ongoing scientific debate (114–122). However, the rigorous pro-inflammatory signaling of Müller cells paired to potential activation of CD4+ and CD8+ T cells in the context of DR should be elaborated in further studies. Towards this goal it should be addressed, whether Müller cells are sufficient to stimulate alloreactive naïve T cells or memory T cells (108).

Furthermore, our proteomic analysis revealed significantly higher abundance of the costimulatory molecule CD40 in pRMG after stimulation with IFNγ. Intriguingly, CD40 has been shown to play a vital role in DR pathogenesis (123–125). A study with transgenic mice, which express CD40 only in Müller cells, demonstrated that the upregulation of CD40 in Müller cells under diabetic conditions is sufficient for thriving leukostasis and degeneration of capillaries, thus promoting retinal inflammation and neurodegeneration in streptozotocin induced DR (124). Expression of CD40 has also been demonstrated in primary human Müller cells and its ligation with CD154 resulted in increased expression of ICAM-1 in Müller cells (123). Furthermore, it was shown that activation of Müller cells through CD40 leads to an enhanced production of TNFα and IL-1β in bystander microglia (124). Since several studies revealed an association between elevated levels of IFNγ and DR, and stimulation of CNS astrocytes led to expression of CD40, IFNγ in DR might lead to the upregulation of CD40 in Müller cells and thus promote chronic inflammation (126–129). Here, we provide new insight in CD40 signaling in Müller cells, as IFNγ increased the expression of CD40, whereas TGFβ2, TGFβ3, TNFα and VEGF resulted in lower abundance of CD40 in pRMG. Future studies are needed to further investigate this issue, as the CD40 signaling pathway constitutes a possibility for therapeutical intervention (130). In contrast to the porcine dataset, we could not identify CD40 in the MIO-M1 cells. As CD40 expression has already been shown in primary human Müller cells, our result might be due to the dedifferentiation of immortalized cells in culture (123).

Oxidative stress and reactive oxygen species (ROS) are known to play a central role during the pathogenesis of DR (131). Rat-derived Müller cells under hyperglycemic conditions developed mitochondrial dysfunction and oxidative stress, causing swelling and eventually apoptosis of the cells (132, 133). Mitochondrial dysfunction can lead to ROS production, which then promotes inflammatory response by activation of NF-κB and release of pro-inflammatory cytokines (134, 135). Our analysis revealed that proteins associated with mitochondrial dysfunction were enriched after treatment of pRMG with all tested cytokines. Furthermore, two significantly enriched pathways in our data sets are associated with reactive oxygen species, namely “NRF2 mediated Oxidative Stress Response” and “Production of Nitric Oxide and Reactive Oxygen Species in Macrophages”. Intriguingly, Müller cells have previously been found to regulate the ROS levels via Nrf2 and to be more resistant to ROS formation compared to photoreceptor cells or bipolar cells (41, 136). In line with this, we showed that treatment with IL-4, TGFβ2, TGFβ3, TNFα and VEGF inhibited death receptor signaling pRMG. Phagocytic cells often produce ROS to protect themselves from pathogens (137). Furthermore, macrophages stabilize cytosolic Nrf2 to be more resistant against ROS (138). Since Müller cells have been shown to by phagocytic, we propose that induction of ROS in these cells also serves as a defense mechanism (109–111).

In this study, we used both primary porcine Müller cells and the spontaneously immortalized human Müller cell line MIO-M1 that was established and described in the Moorfields Institute of Ophthalmology 2002 (39). Although MIO-M1 cells largely retain the phenotypic and functional characteristics of Müller cells *in vitro*, previous studies documented some characteristics of neuronal stem cells and expression of postmitotic neuronal cell markers in MIO-M1 cells (39, 139, 140). Furthermore, it has been shown that Müller cells tend to adapt to cell culture conditions by reduced secretion of neurotrophic factors (141–143). Our analysis revealed many proteins commonly regulated in pRMG and MIO-M1 cells, as well as pathways similarly enriched and regulated in both cells. However, we also observed effects specific for either MIO-M1 cells or pRMG. Thus, for example our IPA revealed that MIO-M1 cells decrease the activity of the significantly enriched canonical pathway oxidative phosphorylation, while it is amongst the most induced pathways of pRMG following treatment with the majority of tested cytokines. While these differences might originate from different adaption of MIO-M1 cells and pRMG to the cell culture conditions or the immortalization of MIO-M1 cells, it might as well be possible that they originate from differences in human and porcine Müller cells. Further studies should elaborate similarities and differences between freshly prepared primary human and porcine Müller cells.

Taken together, our in-depth proteomic profiling of MIO-M1 cells and pRMG revealed the capacity of Müller cells to react in a differentiated manner upon treatment with various growth factors and cytokines. Furthermore, we demonstrated a primarily pro-inflammatory phenotype of Müller cells, as they secreted a variety of pro-inflammatory cytokines, complement components and upregulated proteins associated with antigen processing and presentation, suggesting a function as atypical antigen-presenting cell in the course of retinal inflammation. Furthermore, we observed enrichment of proteins connected to mitochondrial dysfunction, as well as proteins related to the formation and maturation of phagosomes. Thus, Müller cells are capable of modulating immune responses in the retina, and may significantly contribute to chronic inflammation during DR.

## 6 Author Contributions

Conceptualization, S.M.H. and C.A.D.; Formal analysis, S.M.H.; Funding acquisition, S.M.H. and C.A.D.; Investigation, A.S., L.L., C.A.D., and S.M.H.; Project administration, S.M.H.; Supervision, S.M.H.; Visualization, A.S.; Writing - original draft, A.S., L.L., A.G., and D.P.; Writing - review and editing, C.A.D. and S.M.H. All authors have read and agreed to the published version of the manuscript.

## 8 Acknowledgments

The authors would like to thank Adrian Sandbiller for providing porcine eye samples as well as Juliane Merl-Pham for critical discussions.

## 9 Data Availability Statement

All data are contained within the article or in supplementary materials.

