## Supplementary Figures for "Proteomic phenotyping of stimulated Müller cells uncovers profound pro-inflammatory signaling and antigen-presenting capacity"

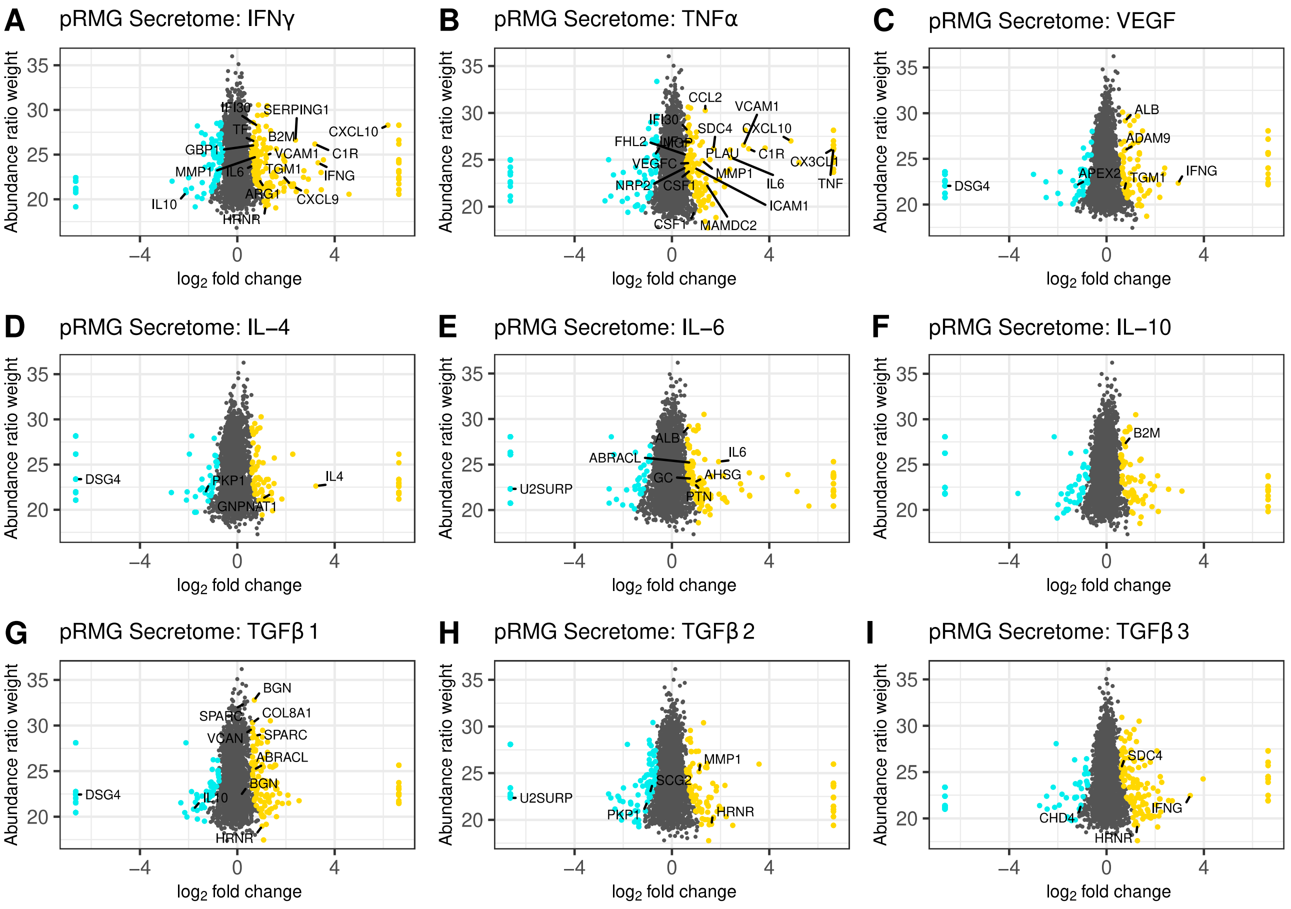


**Supplementary Figure S1.** Scatterplot of all identified proteins from the pRMG secretomes after treatment with the indicated cytokines for 24 h (**A-I**). Proteins with significant changes in their abundance (±$\log_2(1.5)$ fold expression, corrected p-value ≤0.05) were colored, with upregulated proteins being depicted as yellow dots, while down-regulated proteins are colored cyan. Proteins with significantly altered abundance in both, MIO-M1 and pRMG secretomes, are labeled with their gene symbol. Keratins were excluded.

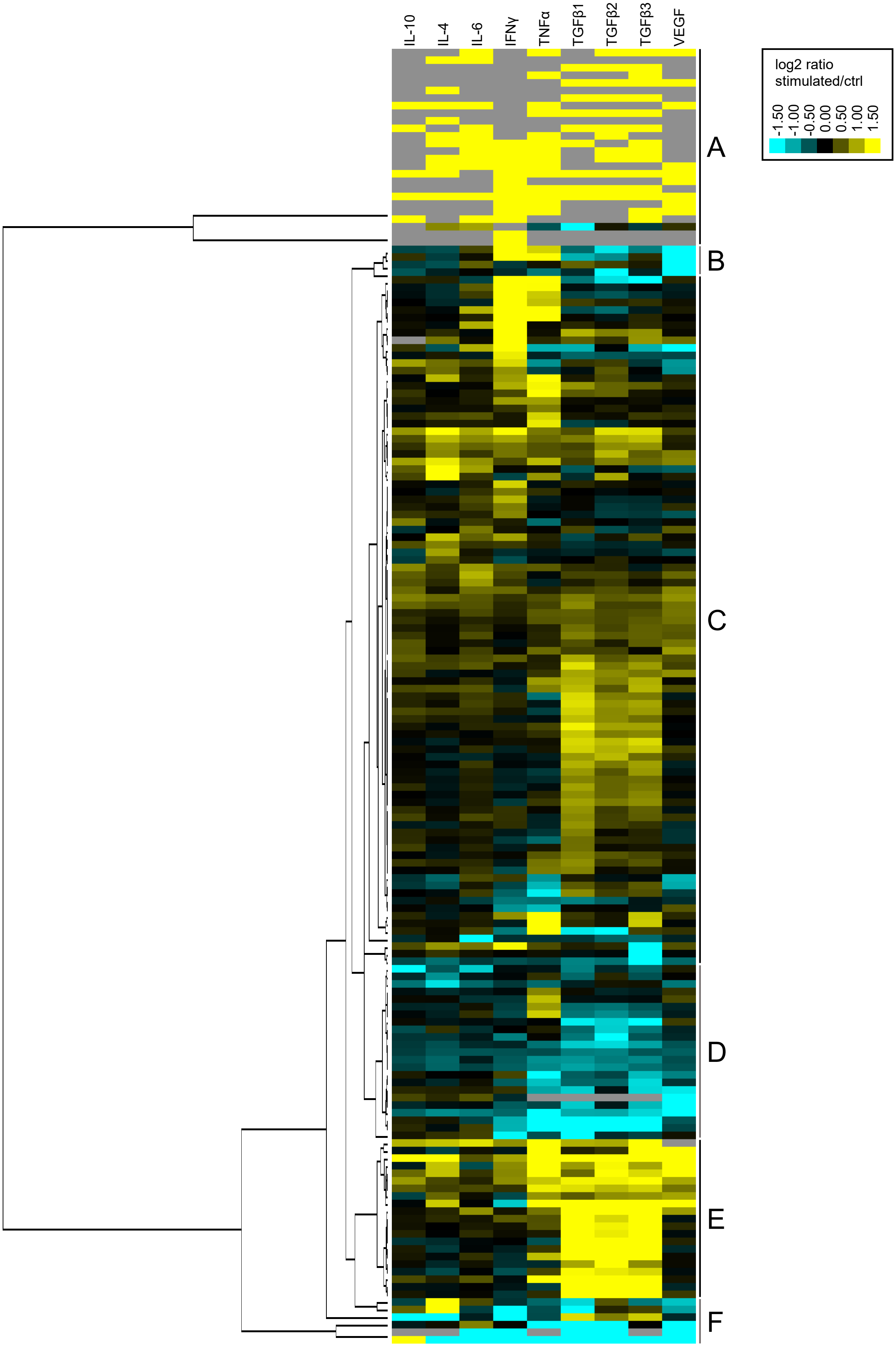
IL-10
IL-4
IL-6
IFNγ
TNFα
TGFβ1
TGFβ2
TGFβ3
VEGF
log2 ratio stimulated/ctrl
-1.50
-1.00
-0.50
0.00
0.50
1.00
1.50
A
B
C
D
E
F

**Supplementary Figure S2.** Heatmap of hierarchical cluster analysis with a dendrogram of proteins secreted by MIO-M1 cells after treatment with various cytokines. Identified proteins were filtered for extracellular proteins with significant changes in their expression (±$\log_2(1.5)$ fold expression, corrected p-value $\leq 0.05$). Down-regulated proteins are presented in cyan, while up-regulated proteins are depicted yellow for the respective treatments. Gray squares represent proteins that were neither identified in the untreated control, nor in the respective treatment. The heatmap was generated on the basis of the log2 fold change of the respective proteins. Clusters A-F were defined using the branches of the dendrogram.

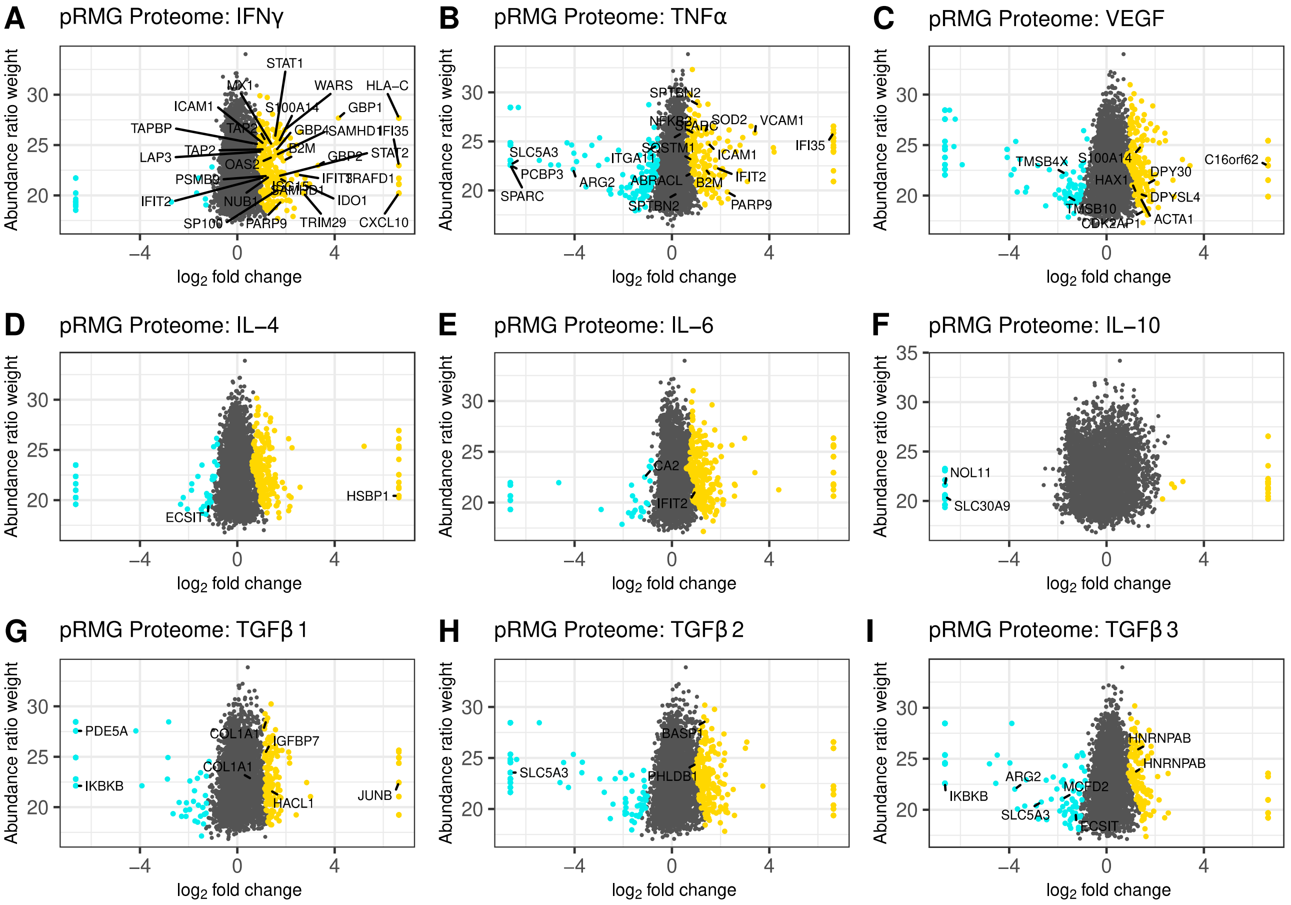


**Supplementary Figure S3.** Scatterplot of all identified proteins from pRMG lysates after treatment with the indicated cytokines for 24 h (**A-I**). Proteins with significant changes in their abundance ($\pm\log_2(1.5)$ fold expression, corrected p-value $\leq 0.05$) were colored, with upregulated proteins being depicted as yellow dots, while down-regulated proteins are colored cyan. Proteins with significantly altered abundance in both, MIO-M1 and pRMG lysates, are labeled with their gene symbol. Keratins were excluded.

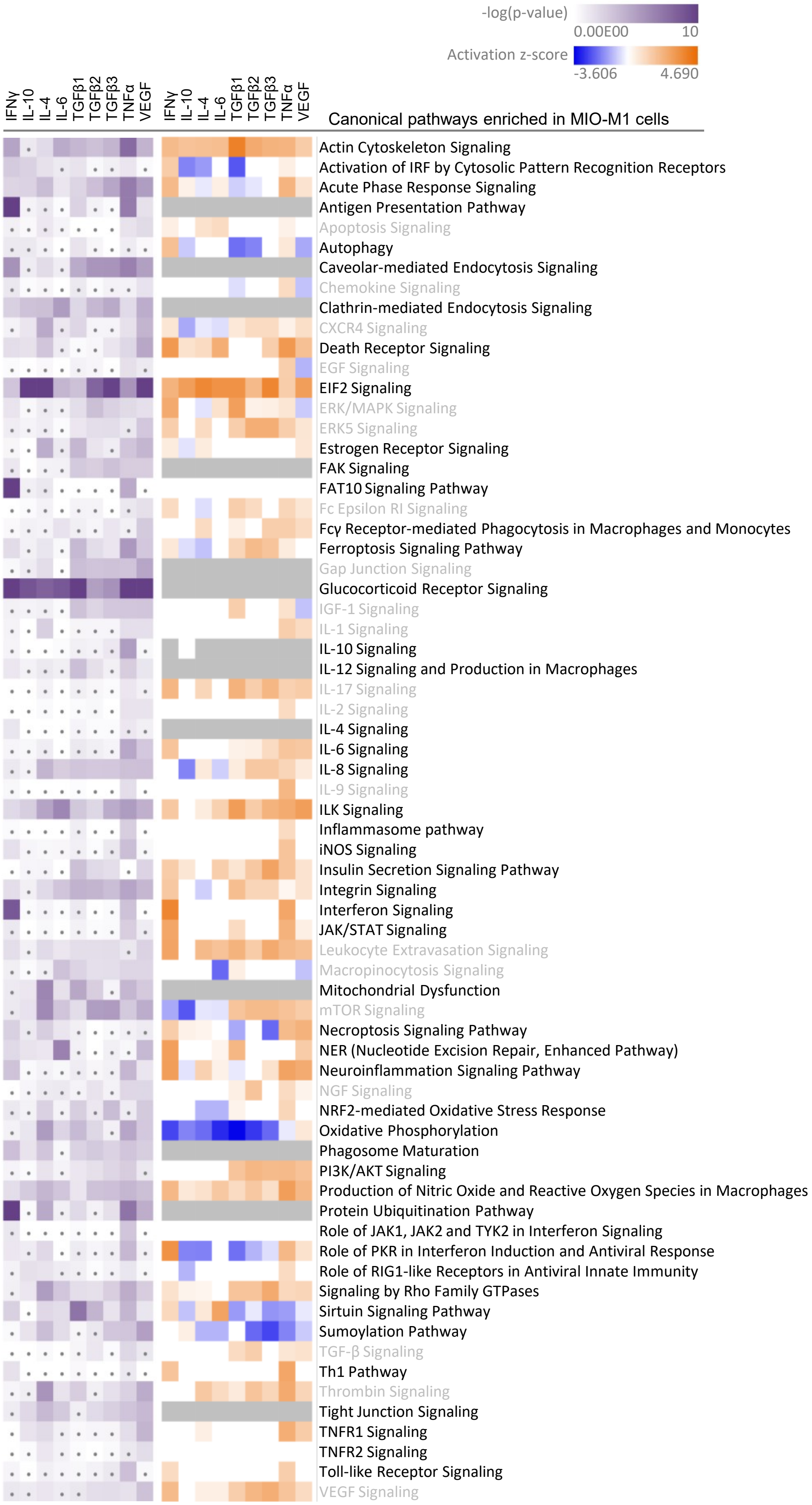
-log(p-value)
0.00E00
10
Activation z-score
-3.606
4.690
IFNγ
IL-10
IL-4
IL-6
TGFβ1
TGFβ2
TGFβ3
TNFα
VEGF
IFNγ
IL-10
IL-4
IL-6
TGFβ1
TGFβ2
TGFβ3
TNFα
VEGF
Canonical pathways enriched in MIO-M1 cells
Actin Cytoskeleton Signaling
Activation of IRF by Cytosolic Pattern Recognition Receptors
Acute Phase Response Signaling
Antigen Presentation Pathway
Apoptosis Signaling
Autophagy
Caveolar-mediated Endocytosis Signaling
Chemokine Signaling
Clathrin-mediated Endocytosis Signaling
CXCR4 Signaling
Death Receptor Signaling
EGF Signaling
EIF2 Signaling
ERK/MAPK Signaling
ERK5 Signaling
Estrogen Receptor Signaling
FAK Signaling
FAT10 Signaling Pathway
Fc Epsilon RI Signaling
Fcγ Receptor-mediated Phagocytosis in Macrophages and Monocytes
Ferroptosis Signaling Pathway
Gap Junction Signaling
Glucocorticoid Receptor Signaling
IGF-1 Signaling
IL-1 Signaling
IL-10 Signaling
IL-12 Signaling and Production in Macrophages
IL-17 Signaling
IL-2 Signaling
IL-4 Signaling
IL-6 Signaling
IL-8 Signaling
IL-9 Signaling
ILK Signaling
Inflammasome pathway
iNOS Signaling
Insulin Secretion Signaling Pathway
Integrin Signaling
Interferon Signaling
JAK/STAT Signaling
Leukocyte Extravasation Signaling
Macropinocytosis Signaling
Mitochondrial Dysfunction
mTOR Signaling
Necroptosis Signaling Pathway
NER (Nucleotide Excision Repair, Enhanced Pathway)
Neuroinflammation Signaling Pathway
NGF Signaling
NRF2-mediated Oxidative Stress Response
Oxidative Phosphorylation
Phagosome Maturation
PI3K/AKT Signaling
Production of Nitric Oxide and Reactive Oxygen Species in Macrophages
Protein Ubiquitination Pathway
Role of JAK1, JAK2 and TYK2 in Interferon Signaling
Role of PKR in Interferon Induction and Antiviral Response
Role of RIG1-like Receptors in Antiviral Innate Immunity
Signaling by Rho Family GTPases
Sirtuin Signaling Pathway
Sumoylation Pathway
TGF-β Signaling
Th1 Pathway
Thrombin Signaling
Tight Junction Signaling
TNFR1 Signaling
TNFR2 Signaling
Toll-like Receptor Signaling
VEGF Signaling

**Supplementary Figure S4.** A comparative IPA analysis with the significantly regulated proteins identified in the MIO-M1 lysates after stimulation with the indicated cytokines was performed. Canonical pathways related to signaling, cell death, immune system processes and the cellular redox state were selected. Pathways with significant enrichment of genes after stimulation with at least one cytokine are presented. Significance of the gene enrichment for each pathway and treatment is indicated by purple squares in the left array. Treatments that did not meet the significance threshold (p-value $\leq 0.05$) are marked with a dot. The z-score is indicated on the right array and represents a prediction of activation (orange) or inhibition (blue) of the pathway. Gray squares mark treatments where the activation state of a pathway could not be calculated. Pathways with significant enrichment in MIO-M1 cells that are not enriched in pRMG cells are labeled gray.
